# Impact of Point Mutation on Shiga-like Toxin 1: A Molecular Dynamics Simulation Study

**DOI:** 10.1101/2023.06.07.544092

**Authors:** Nisat Tabassum, Bristi Rani Paul, Md. Saddam, Md Mostofa Uddin Helal, Susanta Paul

## Abstract

The causative agent of gastroenteritis is Shiga toxin, which belongs to a functionally and structurally associated protein family despite each individual having a unique amino acid sequence. After entering the ER lumen and relocating the toxic domain to the cytoplasm, they alter the large subunit of rRNA, preventing protein synthesis and ribosomal damage. Shiga-like toxin-1 (SLT-1) subunit B targets glycolipid receptor Gb3, which plays a significant role in cytotoxicity. Though the mutational effect on subunit B is important for cytotoxicity study, we lack better understanding. Our present study targets the mutational impact of glycine protein at their 62^th^ amino acid sequence of subunit B. For example, how it can alter the receptor-binding capacity and virulence. We used in silico method with GROMACS software suite (version 5.2, 2020.1) on Google Colab for a 100ns (100,000ps) simulation period and UCSF Chimera software for visualizing mutant and wild-type structure similarities. Surprisingly, RMSD, RMSF, and Rg trajectories from the simulation analysis indicated a more stable and compact mutant structure than the wild type. Principle component analysis (PCA) and SASA were visualized for the entire 100ns, which pointed towards homogeneity between both structures and more solvent accessibility in the mutant structure. This mutation may elevate receptor-binding and virulence capacity. Moreover, this finding can offer a better insight for future vaccine production.

## 1. Introduction

Shiga toxins are mainly found in bacteria *S. dysenteriae*, various serotypes of *Escherichia coli* (STEC), including O157:H7, O104:H4, and Enterobacteria phage H19B(Menge 2020); which plays a major role as a causative agent in severe gastroenteritis and creates disease burden in immunosuppressed patients, infants and elderly population (Thomas et al. 2019). Shiga-like toxins have different isoforms i.e. Stx, Stx1, Stx2, etc. Stx1 and Stx are identical in nature. However, Stx2 isoforms differ immunologically, and approximately 60% similarity to the Stx sequence has been observed. Although each Stx isoform has a unique amino acid sequence, they all have homologous toxin structure and mode of action (Golshani, Oloomi, and Bouzari 2017). Shiga-like toxin 1 (from the Stx1 gene) is a type-II ribosome-inactivating protein; which broadly reside within Shiga toxin family producing functional and structural exotoxins. They enter the endoplasmic reticulum (ER) lumen and create cytotoxicity by relocating their toxic domain to the cytoplasm. These proteins catalytically alter the large subunit of rRNA, preventing protein synthesis and irreversible damage to the ribosome (Basu and Tumer 2015).

Shiga-like toxin 1 contains a single A subunit and pentamers of B subunits (7.7KDa each); which aid in the recognition and binding of holotoxin to cellular globotriaosylceramide Gb_3_ receptor or CD77. Glycolipid receptor Gb3 plays a major role in cytotoxic specificity (Chan and Ng 2016). A trisaccharide receptor analog of Gb3 was used to identify the crystal structure of the Stx1 B subunit in 1998. Three trisaccharide-binding sites were discovered in this study for each B fragment monomer. Any introduction of mutations in the genes producing the Shiga toxin B subunit can have a major impact on its structure, dynamics, and virulence. Moreover, amino acid alteration occurs due to point mutation within the genome; leading to remodeling of stability, binding affinity, and overall toxin-receptor complex behavior(Menge 2020). According to the mutation study, the B subunit has 3 functional sites. Site 1 and 2 play a crucial role in cytotoxicity and mediate high-affinity receptor binding. In addition, site 3 facilitates the identification of additional low-affinity Gb3 epitope (Johannes and Römer 2010).

An in-silico study provides opportunities to study protein dynamics in prediction level. The wild type Shiga-like toxin 1 subunit B is synthesize from stxB gene, which contains 89 amino acids and the Uniport ID is P69179. We aim to elucidate the structural and dynamic changes induced by the mutation and provide valuable insights into the functional consequences of this mutation. The molecular dynamics comparison between (G62T) wild type protein complex and (T62G) mutant protein complex (1CQF) will shed a light on the effect of mutation; as well as corresponding molecular mechanisms underlying disease and target therapeutics development.

## 2. Materials and Methods

### 2.1. Structure Preparation

Model organism Enterobacteria phage H19B gene stxB synthesizes the primary sequence of Shiga-like toxin 1 subunit B protein; which was retrieved from the UniProtKB database in FASTA format and the UniProtKB identifier was P69179. STXB_BPH19. In addition, Expasy’s Prot param server was used to get the physicochemical parameters of protein Shiga-like toxin 1 subunit B. Mutant structure 1CQF was retrieved from RCSB PDB and the DOI is: https://doi.org/10.2210/pdb1CQF/pdb. We also examined the mutant 1CQF protein generated by the UCSF Chimera (Anwar and Choi 2017).

### 2.2. Molecular Dynamics Simulation

A 100 ns (100, 000 ps) molecular dynamics (MD) simulation was conducted in order to assess the stability as well as the consistency of the predicted structure of shiga toxin B. This study was conducted on GROMACS 5.2 (2020.1) (Abraham et al. 2015) on an on google colab framework. For shiga toxin B structure, MD simulation of mutated protein was carried out. This simulation not only performed to compare the mutated structure trajectory data but also to anticipate the shared similarities between trajectory analysis of both wild-type and mutated structure prediction. The protein topology force field was developed utilizing all-atom Optimized Potentials for Liquid Simulations (OPLS-AA) (Robertson, Tirado-Rives, and Jorgensen 2015). In particular, the non-bonded interaction parameters calculated by OPLS-AA and OPLS-AA/L show positive findings (Shirts et al. 2003; Tzanov, Cuendet, and Tuckerman 2014). Besides, both the wild type and mutated protein structure was solvated using a general equilibrated 3-point solvent model named simple point charge (SPC) water model spc216. A 1.50 nm cubic simulation box was constructed around the projected model and solvated using an SPC water model. 4 cl^-^ ions were added to neutralize the positive charge. In order to eliminate the edge effect, periodic boundary requirements were imposed in every direction. Both the mutant and the wild type protein structures were used in the system’s energy minimization, which was done using the steepest descent algorithm with a maximum step size of 50,000 and a tolerance of 1000 kJ mol^-1^ nm^-1^. Following system minimization, it was equilibrated for 100 ps at both the isothermal-isobaric ensemble (NPT) and the canonical ensemble (NVT). For establishing long-range electrostatic interactions with a PME order, the Particle Mesh Ewald (PME) summation was utilized. Restricting hydrogen-atom bonds and water-molecule geometry, the Linear Constraint Solver (LINCS) algorithm and SETTLE technique were employed (Hess et al. 1997; Miyamoto and Kollman 1992). The Parrinello-Rahman method modulated a constant level of pressure at 1 atm (1.01325 bar); while at the same time, temperature regulation at 300K was achieved using V-rescale weak coupling method. For the 100 ns MD run production with no constraints, the LINCS algorithm and a 2fs (fs) integration stage was utilized (Hess et al. 1997). PME method was utilized for Lennard-Jones and Coulombic interactions.

### 2.3. Molecular Dynamics Analysis

A widely used method for deriving functionally important collective movements from a molecular dynamics (MD) trajectory is principal component analysis. We used previously known methodologies to conduct an examination of the essential dynamics (Anwar and Choi 2017; Spellmon et al. 2015; Wolf and Kirschner 2013). Also, we employed MDAnalysis and MDTra’s atom selection and associated algorithms to evaluate the C-alpha backbone trajectory as well as slice the trajectory over the entire 100ns (McGibbon et al. 2015; Michaud-Agrawal et al. 2011). The RMSD distance matrices and hierarchical clustering techniques were also employed to group the MD simulation trajectory of shiga-like toxin 1 subunit B and the mutated structure 1CQF, using the MDTraj Python module. Every pairwise RMSDs between conformations are computed, and the result is a dendrogram with RMSD average linkage hierarchical clustering. Lastly, by mapping the simulation data into the reduced dimensional space of both the modelled and mutated structures, we created a two-component PCA model to calculate the main components of the total 100 ns while taking into account the alpha carbon chain. The alignment-dependent input PCA leverages Cartesian coordinates. In comparison, we calculated the alignment independent pairwise distance PCA between every atom in each frame of the modeled and mutated alpha-carbon chain.

## 3. Result and Discussion

### 3.1. Structural Properties

We used the UniProt database for retrieving the whole structure of the wild-type Shiga toxin-1 B component (UniProt ID: P69179) and RCSB PDB database for mutant protein 1CQF (DOI: https://doi.org/10.2210/pdb1CQF/pdb). Mutation was introduced in the Shiga toxin B subunit’s wild-type structure by changing the residue G62 to T62 using Chimera software program. The amino acid threonine (T) at the 62^th^ position was mutated to glycine (G). This resultant mutant structure 1CQF represents the complex of the mutated Shiga toxin. Once the mutation had been introduced via UCSF Chimera (Pettersen et al. 2004), the combination of the mutant and wild type Shiga toxin was carried out.

### 3.2. Fundamental Dynamics Analysis

Molecular dynamics simulations were performed to investigate the structural dynamics and stability of wild-type (G62T) and mutant (T62G) structures. For the simulation we followed the article of Paul et al., 2022; which used Google Colab’s optimized simulation protocol. The MD simulations were carried out using GROMACS 2020.1 software, utilizing the GPU service on Google Colab to leverage its computational capabilities. The simulations were run for 100 ns to illustrate the behaviour of the protein over an extended period (Paul et al. 2022).

RMSD is a quantitative measure to evaluate the similarity between multiple protein structures. In order to determine the structural stability, the RMSD of the protein backbone was calculated for the wild type (G62T) and mutated (T62G) structures over a 100 ns (100,000 ps) simulation. The RMSD values were obtained by comparing the final structures to their initial conformations. **Figure 1A** displays the maximum and minimum distances of RMSD for the wild and mutant type trajectories, which were approximately 0.34 nm and 0.17 nm respectively. Additionally, six distinct intersections were identified within these trajectories at 10 ns, 30 ns, 45 ns, 65 ns, 90 ns, and 97 ns. In summary, both the wild type and mutant structures reached a stable state around 97 ns, with an RMSD range of 0.17-0.35 nm. These findings suggest that both structures achieved equilibrium and maintained their overall structural integrity throughout the simulation.

**Figure 1.**
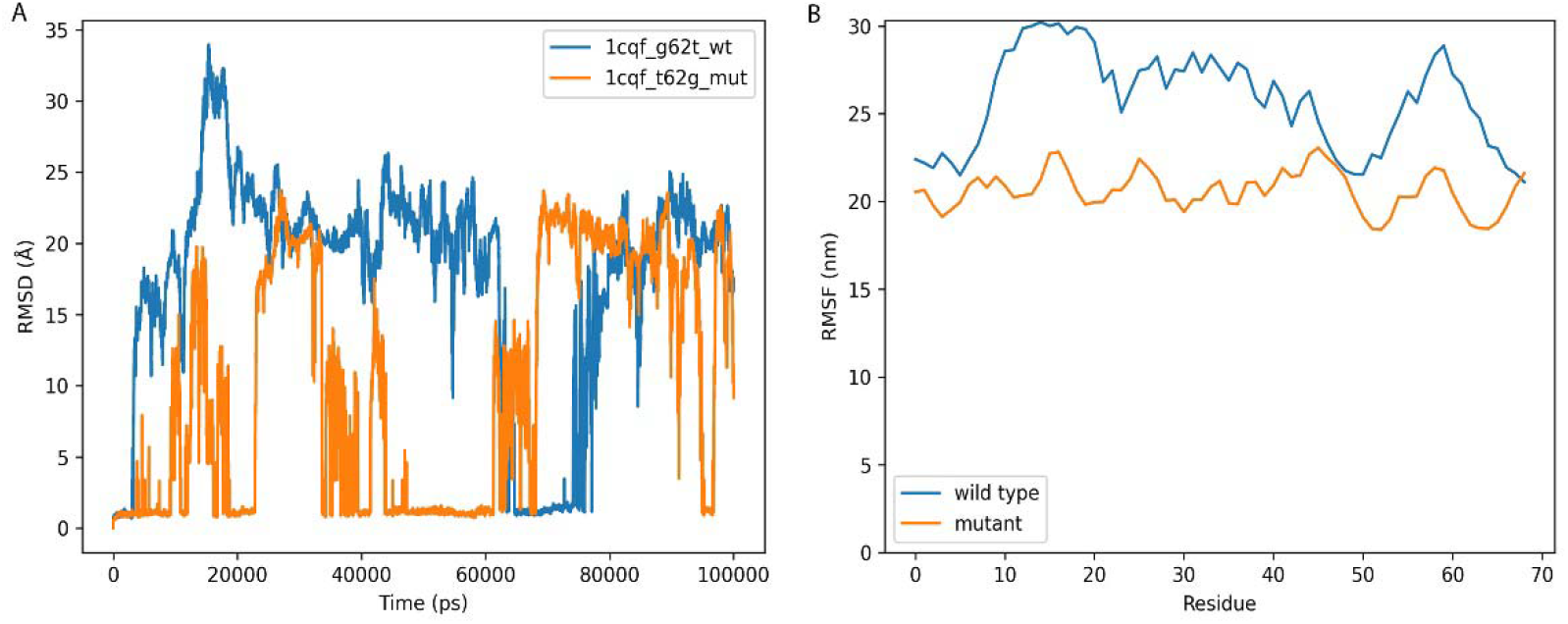
The results of simulation (A) Representing the RMSD of wild (blue) and mutant (orange) type during the entire 100ns. (B) Trajectories of RMSF fluctuation of G62T wild type (blue) and T62G mutant (orange) showing three flexible region. In (A), from little to no deviation both structures started showing variations within few nano seconds. The highest deviation between two trajectories was about 0.34nm, which remained for about 10ns (from 15ns-25ns). The system gets stable in the range of 0.17-0.35 nm at about 97ns by the convergence of the both structure. (B) RMSF data identifies 3 significant deviations between two trajectories of about 10nm, 7nm, 6nm respectively.

The RMSF is an averaged measure of mutant complex by which we predict the displacement of a specific atom, or cluster of atoms, with respect to the wild-type structure. The investigation of the structure’s time-dependent movements are retrieved from the RMSD. **In Figure 1B**, we analyzed the flexibility of the structure; which represents the Backbone RMS Fluctuation (RMSF) of the wild and mutant structure. RMSF data identifies 3 significant deviations of amino acid residues. The first one is between 8-23th residues, where the deviation is from 23-30nm; the second fluctuation is between the 28-45 residue while 17-28nm is the deviation range and finally, from 50-68th residue the deviation ranges from 21-28nm. Comparing the architecture of mutant and wild-type organisms enabled us to evaluate of the effect of the T62G mutation on the complex’s flexibility. From the RMSF value we can clearly visualize that, the mutant type structure is more compact and has less deviation in its trajectory representing its stability. Integration of glycine in the 62th glycine rich loop potentially increased ligand binding capacity in the mutated protein.

In **figure 2**, the radius of gyration analysis was conducted to evaluate the compactness and global structural changes of the protein during the simulation. One of the most important metrics that is frequently used to forecast the structural activity of a macromolecule is the computation of Rg. The result resembles seven different intersections; started at the 15ns, then at 25-35ns, the third on at 45ns, in between 60ns-70ns the fourth convergence appeared, later at 80ns-90ns, 95ns and at last the final convergence was from 98-100ns. This result clearly stated that, the profile of the T62G mutants’ compactness was closely resembled to that of the wild type and showing an analogy in their overall homogeneity.

**Figure 2.**
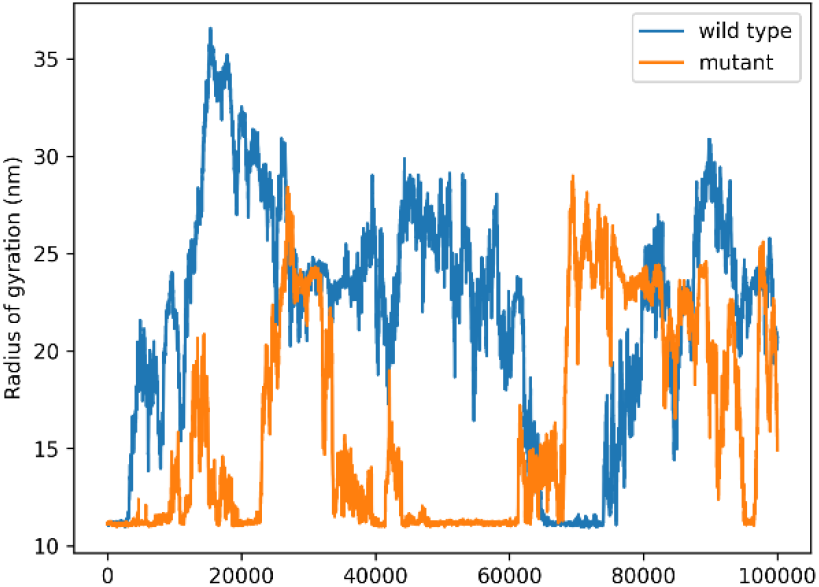
Representing the Radius of gyration of (Rg) of wild type (blue) and mutant (orange) structures. The Rg data shows overall compactness of the wild and mutant structure.

#### 3.3. Essential Dynamics Analysis

Principal component analysis (PCA) was performed to analyze the collective motions and conformational changes in the Shiga toxin wild type (G62T) and mutated (T62G) structures. Pairwise distance PCA is calculated by analyzing the atoms precise location against time. On the other hand, the Cartesian coordinate PCA analysis partially captures the dominating overall motion. The hierarchical distribution of all the clusters is shown via RMSD hierarchical clustering. This distribution was visualized using color coded clusters against a fixed time frame and the color distribution ranged from initial to final stage at 100ns simulation in both Cartesian coordinate PCA and pairwise distance PCA.

Complexes produced in **figure 3, 4 and 5** evaluating the conformational dynamics of the mutant structure (1CQF) in comparison to the wild type structure and the study comprises clustering dendrograms (**figure 3**) and Principal Component Analysis (PCA) plots (**figure 4 & 5**).

**Figure 3.**
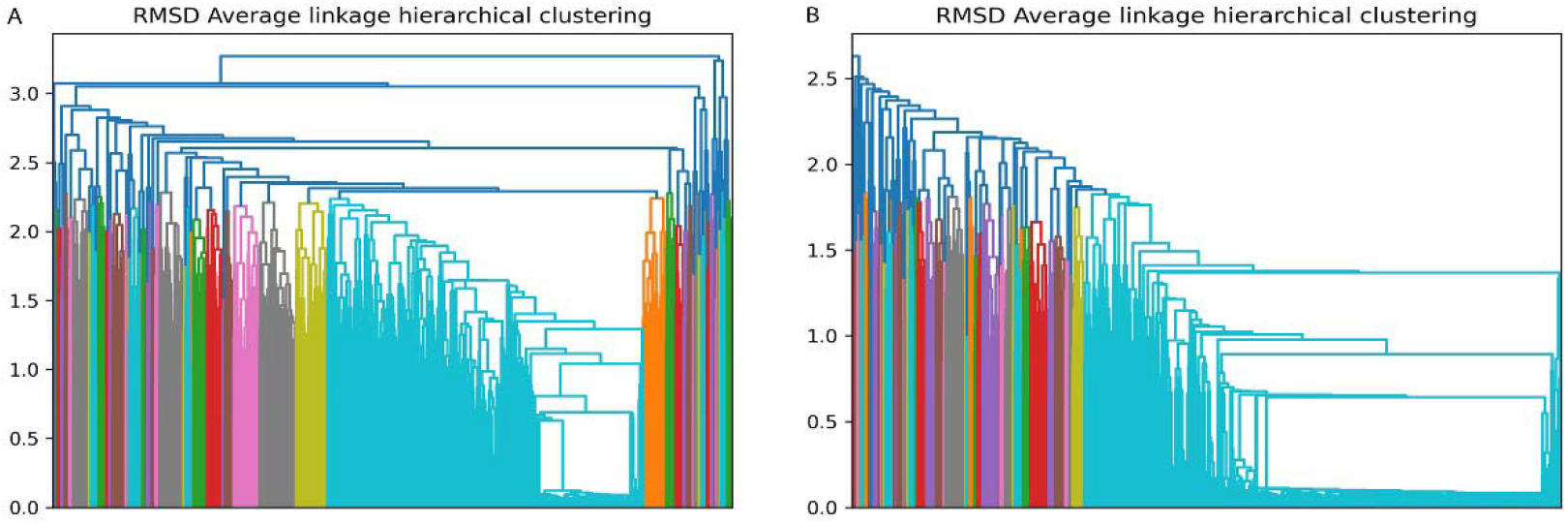
Essential dynamics analysis of (**A**) wild and (**B**) mutant structure. It represents clustering dendrograms of both structures for entire 100ns of simulation period. The hierarchical distribution of all the clusters is shown via RMSD hierarchical clustering.

**Figure 4.**
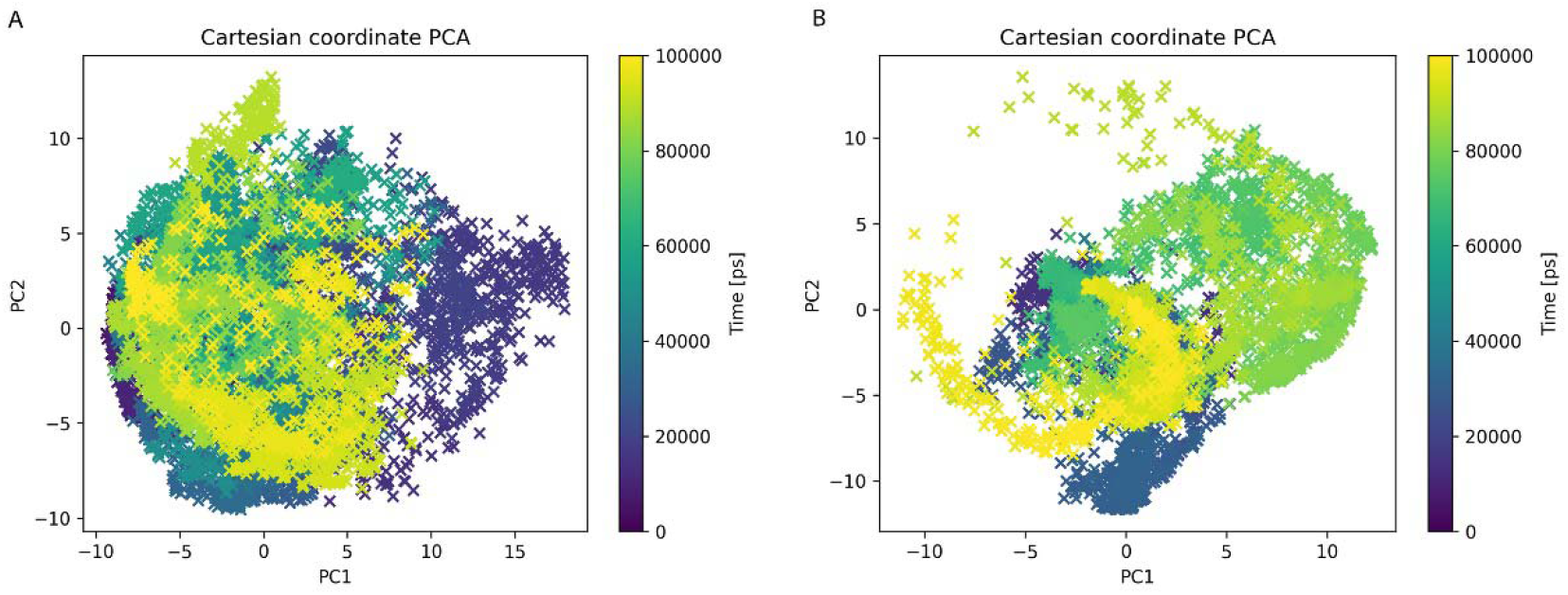
Visualizing Cartesian coordinate (PCA) principal component analysis of wild type (**A**) and mutant (**B**) structure of entire 100ns of simulation period. These distributions were visualized using color coded clusters against a fixed time frame. In (**4A**) wild type structure, we visualized the initial clusters were residing between first and second coordinates and in the final stage they were placed within 3rd and 4th quadrants. Though in (**4B**) mutant, primary clusters were within 2nd and 3rd quadrants; their final appearance was between 3rd and 4th quadrants as well.

**Figure 5.**
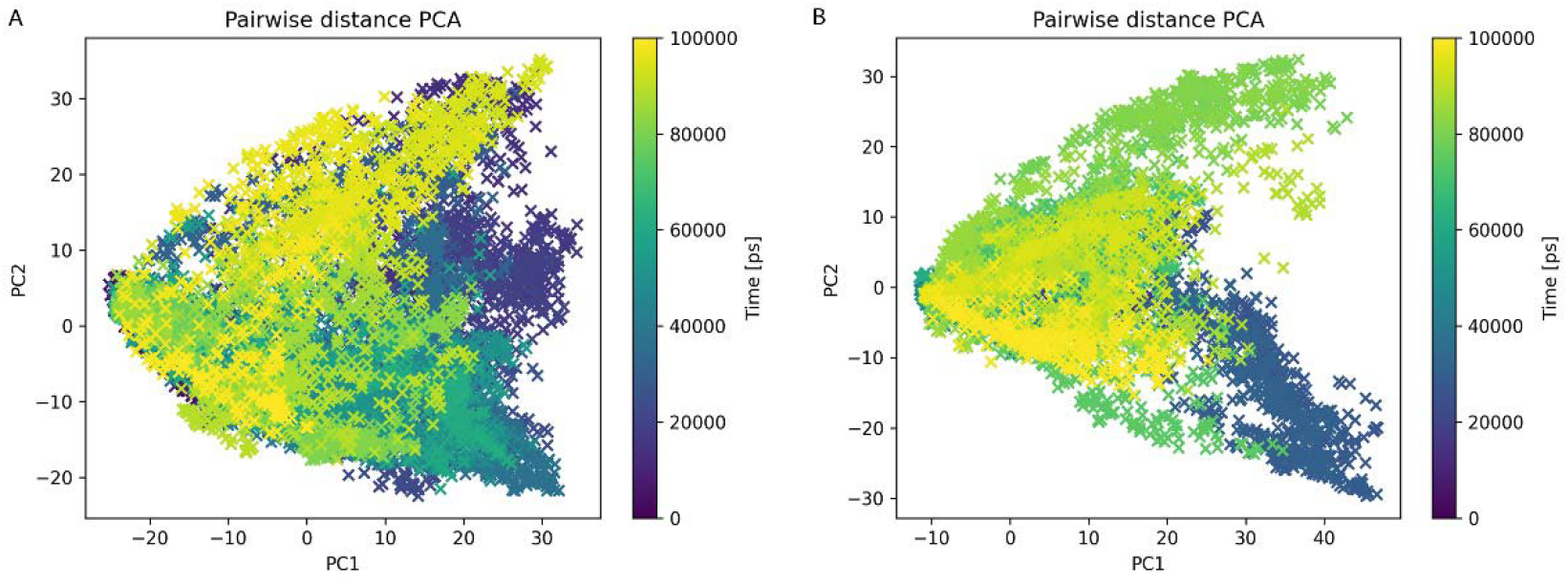
Visualizing pairwise distance (PCA) principal component analysis of wild type (A) and mutant (B) structure of entire 100ns of simulation period. These distributions were visualized using color coded clusters against a fixed time frame. In pairwise distance PCA (**5**), final clusters of wild type (**5A**) were visualized between 2nd and 3rd quadrants. In contrast, mutant type (**5B**) showed their final clusters in 3rd and 4th quadrant.

The Cartesian coordinate PCA plots in f**igures 4A** and **4B** depict the entire 100ns of the whole simulation period for the wild type (**A**) and mutated structure (**B**) against a **2D** graph respectively. In **figure 4A**, we visualized initial clusters were residing between first and second coordinate, in the final stage they were placed within 3rd and 4th quadrant. Though in **figure 4B**, primary clusters were in 2nd and 3rd quadrants; their final appearance was between 3rd and 4th quadrants as well. In pairwise distance PCA, final cluster of wild type was visualized between 2nd and 3rd quadrants. In contrast, mutant type showed their final cluster in 3rd and 4th quadrant.

The solvent-accessible surface area (SASA) was calculated using the Shrake and Rupley algorithm of MdTraj (McGibbon et al. 2015). In **figure 6**, we investigated the total Solvent Accessible Surface Area (SASA) analysis for the wild type (A) and mutated (B) structure; which is necessary for hydrophobic core region analysis for precise understanding of the stability, binding interaction and folding pattern of the protein. The solvent-exposed region was discovered using SASA analysis, which indicates towards conformational change of mutant protein (**B**) in comparison to wild type (**A**) protein.

**Figure 6.**
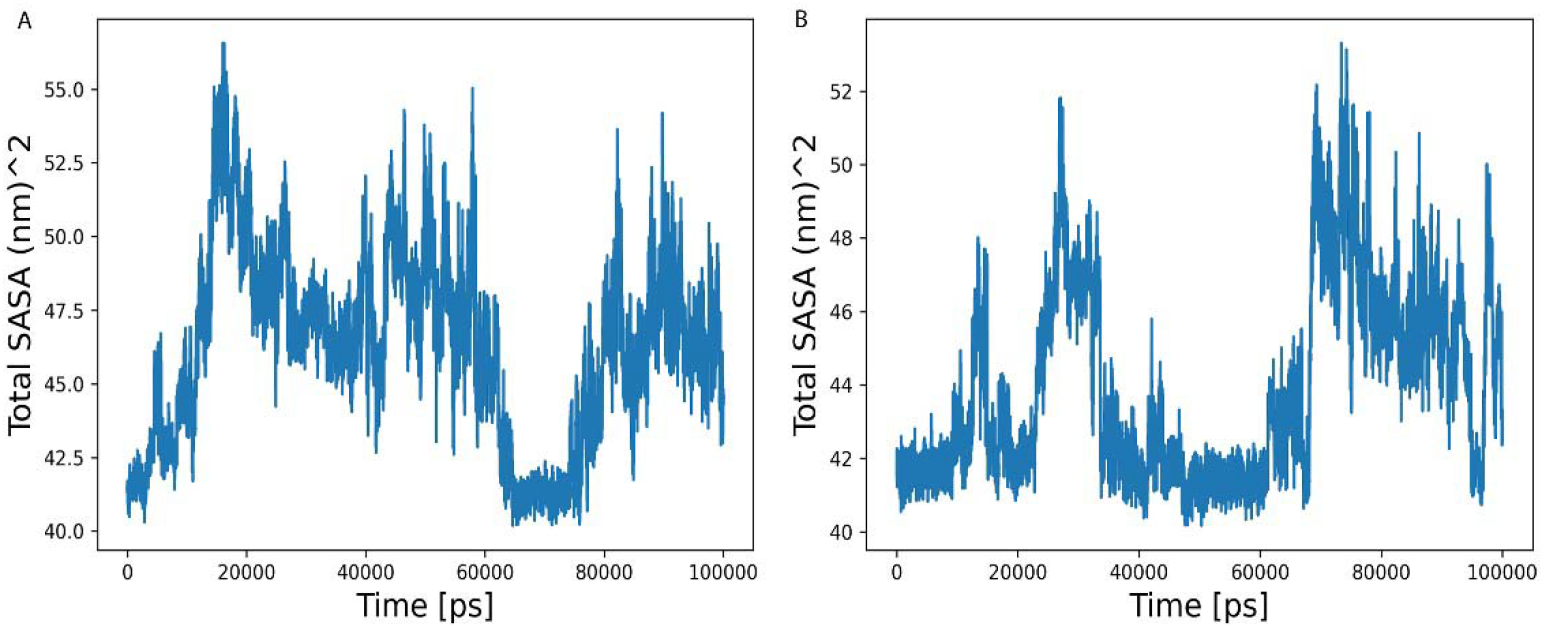
Depicts total SASA value of wild (**A**) and mutant type (**B**) structure of entire 100ns. Where wild type (**6A**) showed highest peak value of about 58 nm^2^ in between 18-20 ns. Mutant type (**6B**) showing the highest value of about 55 nm^2^ at approximately 75 ns.

In **figure 7**, SASA autocorrelation was depicted. The wild type (**A**) variant maintained a constant value of about 1nm^2^ from the 0.10ns-1ns; then a constant downfall until two peaks of about 0.1nm^2^ (40-60ns) and 0.2 nm^2^ (70-90ns) has been observed between 10-100ns. Similar pattern has been observed in case of mutant structure (**B**) with an exception of a single peak of about 0.3 nm^2^ was observed between 40ns-90ns. Despite having functional homogeneity between wild (**A**) and mutant type (**B**) structure we can observe that, mutant structure (**B**) achieved more improved binding capacity and thus increased virulence frequency.

**Figure 7.**
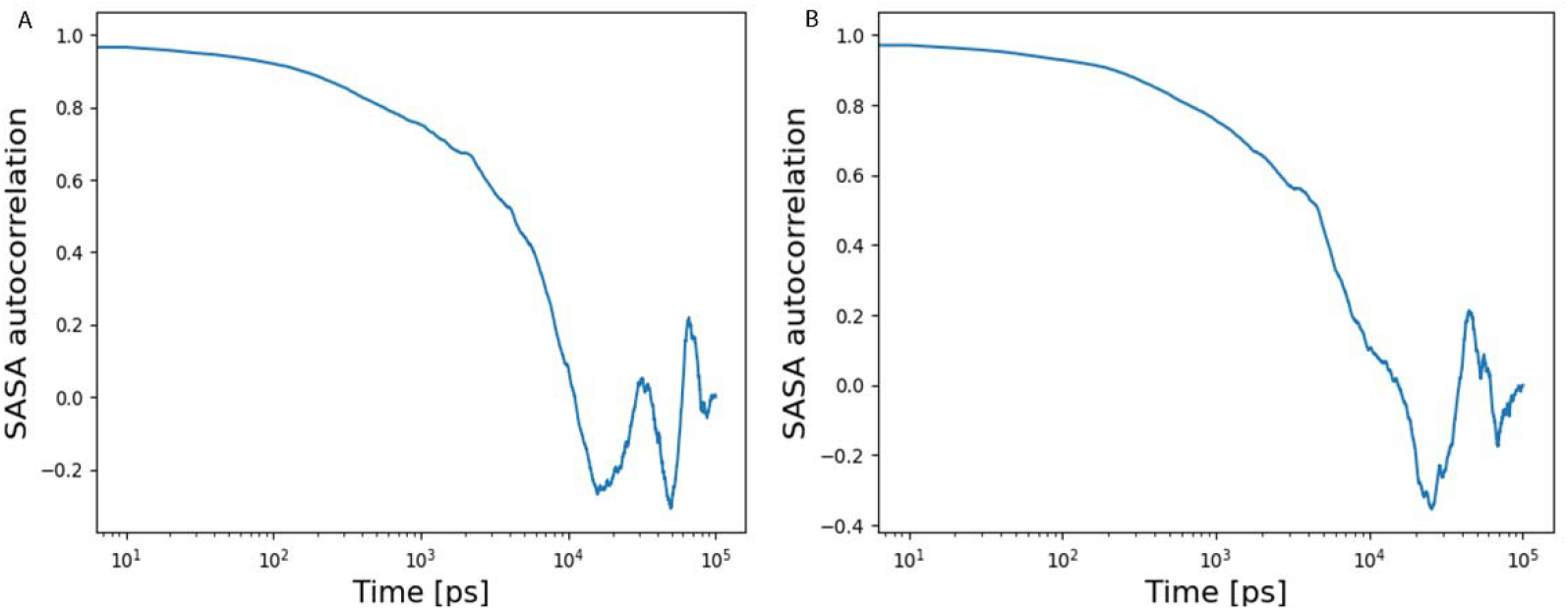
Depicting SASA autocorrelation. The wild type (**7A**) variant maintained a constant value of about 1nm^2^ from the 0.10-1ns, then a constant downfall until two peaks of about 0.1nm^2^ (40-60ns) and 0.2nm^2^ (70-90ns) has been observed between 10-100ns. Similar pattern has been observed in case of mutant structure (**7B**) with an exception of a single peak of about 0.3nm2 between 40ns-90ns.

## 4. Conclusion

From our in silico investigation, we analyzed the mutation induced structural changes, pathogenic pattern and changes in binding affinity. Molecular Dynamics simulation of both wild and mutant Shiga toxin structure predicted RMSD, Radiation of gyration (Rg), RMSF value. After assessing the RMSD value, overall resemblance and homology between wild and mutant type structure was identified. We targeted both of our complexes for their compactness and time dependent movement by RMSF. Our study investigated a more compact (T62G) mutated structure 1CQF with a highest deviation of 28nm in comparison of the wild type structure having a deviation of 38nm. It clearly pointed towards a more compact receptor-binding site along with increased virulence capacity in the mutated protein. The Radius of Gyration (Rg) calculated the structural activity of a protein. Our investigation finds out similarities between protein complexes along with a highly stable mutant protein structure. Along with this, SASA indicates towards conformational changes and improved binding capacity. We obtained a precise understanding of the intramolecular properties of both protein structures and their overall dominating motion; mainly through the pairwise distance and Cartesian coordinate PC. Principle component (PC) analysis clearly stated homogeneity between (T62G) mutant protein and (G62T) wild type protein. In concise, the mutation at 62^th^ amino acid threonine residue to glycine not only increased the receptor-binding capacity but also it could be a stimulant factor for increased virulence capacity in the mutant Shiga-like toxin type-1 subunit B.

## Acknowledgements

We express our gratitude to Shamrat Kumar Paul for generously providing us with simulation and analysis source code.

